# Contrast-enhanced microCT evaluation of degeneration following partial and full width injuries to mouse lumbar intervertebral disc

**DOI:** 10.1101/2022.01.14.476362

**Authors:** Remy E. Walk, Hong Joo Moon, Simon Y. Tang, Munish C. Gupta

## Abstract

**Study Design:** Preclinical animal study

**Objective:** Evaluation of the degenerative progression resulting from either a partial- or full- width injury to the mouse lumbar intervertebral disc (IVD) using contrast-enhanced micro-computed tomography and histological analyses. We utilized a lateral-retroperitoneal surgical approach to access the lumbar IVD, and the injuries to the IVD were induced by either incising one side of the annulus fibrosus or puncturing both sides of the annulus fibrosus. The full-width injury caused dramatic reduction in nucleus pulposus hydration and significant degeneration. A partial-width injury produces localized deterioration around the annulus fibrosus that resulted in local tissue remodeling without gross degeneration to the IVD.

**Methods:** Female C57BL/6J mice of 3-4 months age were used in this study. They were divided into three groups to undergo partial-width, full-width, or sham injuries. The L5/L6 and L6/S1 lumbar IVDs were surgically exposed using a lateral-retroperitoneal approach. The L6/S1 IVDs were injured using either a surgical scalpel (partial-width) or a 33G needle (full-width), with the L5/L6 serving as an internal control. These animals were allowed to recover and then sacrificed at 2-, 4-, or 8-weeks post-surgery. The IVDs were assessed for degeneration using contrast-enhanced microCT (CEµCT) and histological analysis.

**Results:** The high-resolution 3D evaluation of the IVD confirmed that the respective injuries localized within one side of the annulus fibrosus or spanned the full width of the IVD. The full-width injury caused deteriorations in the nucleus pulposus after 2 weeks and progressed to significant degeneration at 8 weeks, while the partial width injury caused localized disruptions that remained limited to the annulus fibrosus.

**Conclusion:** The use of CEµCT revealed distinct IVD degeneration profiles resulting from partial- and full- width injuries. The partial width injury may serve as an alternative for IVD degeneration resulting from localized annulus fibrosus injuries in humans.

## Introduction

Mouse models are often used for the preclinical validation of treatments and therapeutic candidates. Mice offer particular advantages of easy maintenance, year-round breeding, short gestation period, large litters, and inbred tolerance^1^. Furthermore, the ability to manipulate the mouse genome enables mechanistic studies with greater biological precision than larger mammalian models^1^. Moreover, the mouse lumbar intervertebral disc (IVD) provides close geometric and microstructural semblance to the human lumbar IVD compared to other preclinical animal models^2^. While the mouse lumbar IVD exhibits age-related degeneration^3,4^, an injury is often utilized to reproduce the inflammatory conditions and accelerate the IVD’s degenerative cascade^5,6^.

To create a targeted injury to the IVD, surgical exposure of the IVD is required. These commonly involve the posterior-lateral, transperitoneal, and anterior/lateral retroperitoneal surgical access. Of these, the retroperitoneal approach particularly advantageous in that it : (1) does minimal damage to major organs, vessels, and musculature; (2) accesses multiple spinal levels with a relatively small incision; and (3) enables direct visualization of the target tissue under microscopy. Moss et al. demonstrated the efficacy and reproducibility of the retroperitoneal approach in rabbits ^7^. Exposing the lumbar IVD enables the ability to create a targeted, directed injury for the investigation of the subsequent degenerative process ^8–10^. Masuda et al. and Sobajima et al. described the rabbit annulus fibrosus injury model where the injury was confined to the annulus fibrosus that caused slow progressive degeneration of the IVD over 8 weeks. The phenotype and the progressive nature of this model recapitulates the human disease, and may be better suited to explore therapies that leverage prevention or regeneration. In contrast, more damaging approaches that injures both the annulus fibrosus and nucleus pulposus produce rapid and severe course of degeneration ^7,11^. There are several studies describing IVD degeneration using the lumbosacral IVD injury in the mouse model^12–14^. In the mouse IVD where the average disc height is approximately 300-400 microns, an injury by needle puncture produces damage to 55-90% across the height and 15-40% across the width of the IVD^2,11^, representing significant trauma that is atypical in human IVDs. Moreover, these procedures involve damage and injury to the nucleus pulposus, such as with a complete puncture to the IVD, with no possibility to decouple the contributions of the annulus fibrosus and the nucleus pulposus toward the ensuing degenerative cascade. Piazza and coauthors have shown that unilateral (half-width) and bilateral (full-width) injuries in the tail IVD results in unique degenerative trajectories, but this has not been investigated in the mouse lumbar spine. We thus sought to compare the degenerative profiles of IVDs after partial- and full- width injuries in the mouse lumbar spine. In order to evaluate the degeneration of the IVD in a spatially robust manner, we utilized contrast-enhanced microCT^15,16^, in addition to histological grading, to quantify the changes in structure and composition at 2-, 4- and 8-weeks after surgery.

## Materials and Methods

### Animal Preparation

All procedures were performed following Washington University School of Medicine IACUC approval. Female C57BL/6J mice of 3-4 months age were used (BW: 20 – 25 g). They were housed under standard animal husbandry conditions (in a temperature-controlled [21 ± 1°C] room with normal 12-hr light/dark cycles). These animals were divided into three groups: Partial-width injury (PW), full-width injury (FW), and Sham (n=15-18 per group) with all animals undergoing to retroperitoneal surgical exposure of the lumbar IVD. The groups were cross-sectionally evaluated at 2-, 4-, and 8- week post-surgery time points. The injury was delivered to the L6/S1 IVD with the L5/6 IVD used as the internal control.

The partial-width injury mimics a localized injury to the annulus fibrosus, aka annular tear injury, in humans^17^. To evaluate the feasibility of this injury, we first performed the partial-width injury on six animals, and then they were euthanized shortly after recovery from anesthesia. The IVDs were harvested and measured for the thickness of the annulus fibrosus and the depth of the injury using contrast-enhanced microCT (CEµCT) and histological analysis.

The second set of animals were allowed to recover following PW, FW or Sham surgery for 2, 4 or 8 weeks (n = 5-6 per group) and then euthanized, and the IVD tissues were assessed for degeneration with CEµCT and histological analyses.

### Retroperitoneal approach to the intervertebral disc procedure

Mice were anesthetized with isoflurane gas in oxygen via a facemask (3–4% induction and 2–2.5% maintenance at 1 L/min flow rate; Highland Medical Equipment) and were given a preoperative intradermal injection of lidocaine (7 mg/kg; Hospira, Inc). The left flank was then shaved from the ventral to the dorsal midlines, and the skin was sterilized. The skin was prepared for aseptic surgery via washing with 70% ethanol and povidone iodine. Under microscopic guidance, retroperitoneal dissection was used to expose the lateral aspect of the spine with a Penfield dissector. The pelvis could be rotated posteriorly to expose a broad working space while holding the pelvic bone (Figure 2A). The Penfield dissector was pulled out the peritoneal wall to expose the psoas muscle and to protect the abdominal organs (Figure 2B). The psoas muscle was stripped posteriorly to anteriorly by initially scraping out tissues attached to the anterior surface of the pelvis (Figure 2C). The spinal column and the intervertebral disc were exposed from the ventral midline to the posterior margin of the disc at the L5/6 and L6/S1 levels (Figure 2D). Since the superior margin of the pelvic bone indicates the L6 vertebral body (Figure 2E), the location of the L5/6 and L6/S1 IVD could be confirmed (Figure 2D and 2E). Extension to the cranial space beyond the L5/6 was also easily achieved by blunt dissection. Additional details and images on the surgical procedure can be found in Supplemental methods, Supplemental Figure S1, and Supplemental video.

### Partial-width and full-width injuries

For the partial-width injury, a distance of 0.3 mm from the end of the No. 11-scalpel blade tip was measured and marked with a micro-caliper under a microscope (Figure 1). The distance of 0.3 mm was determined from our preliminary studies. The edge of the blade was allowed to insert into the IVD until the 0.3 mm marking was no long visible under the microscope. The injury site was closely observed under the microscope to confirm that there is no leakage of the NP.

**Figure 1.**
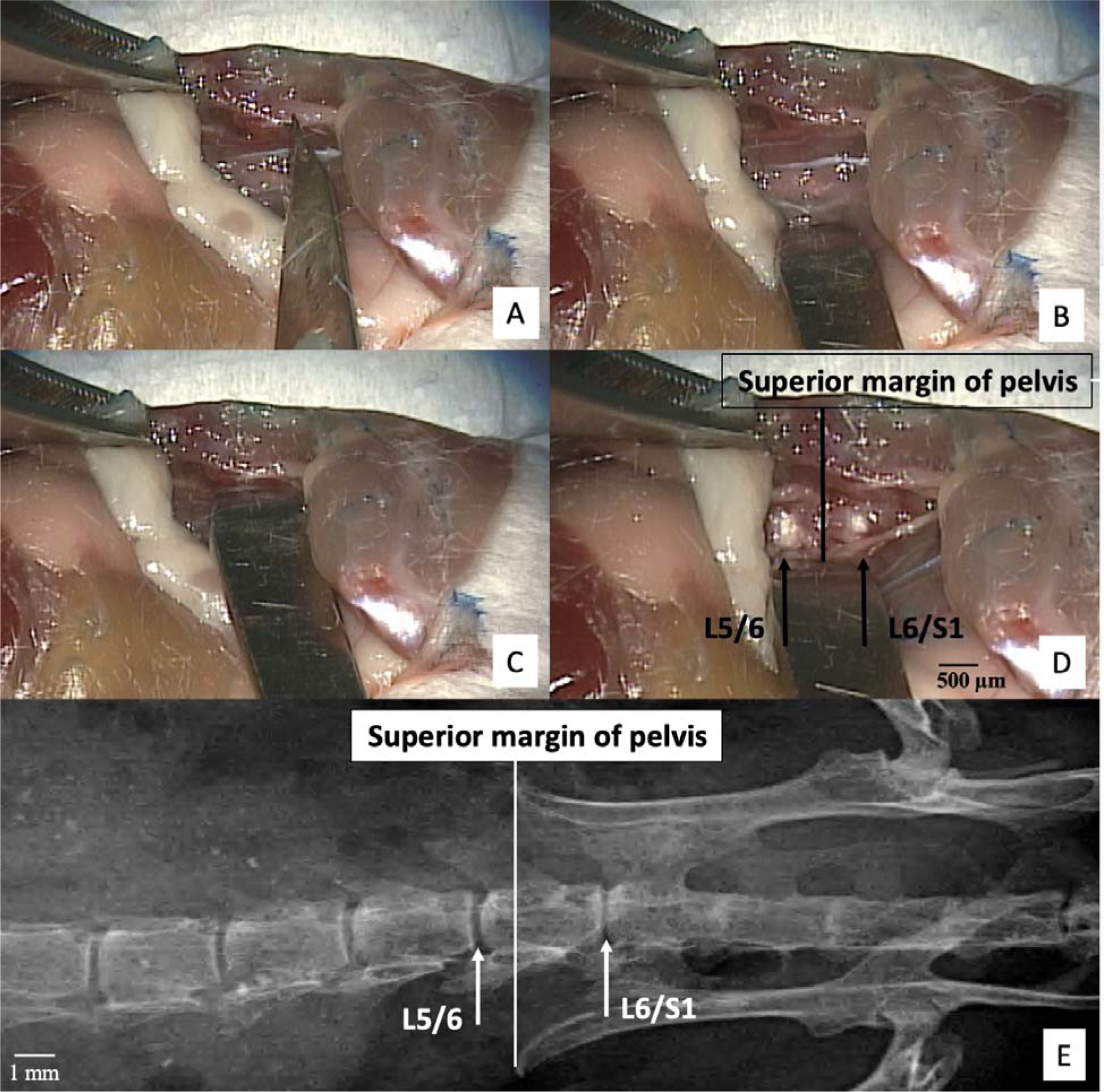
Retroperitoneal dissection to expose the lumbosacral intervertebral disc using microscopic guidance **(A)** The No. 11-scalpel blade indicates the left pelvic bone and forceps grip the gluteus muscles. (**B)** The pelvis can be rotated posteriorly to increase working space by raising the gluteus muscles and rotating the left pelvic bone. The abdominal wall and peritoneum are retracted anteriorly by a Penfield dissector to expose the psoas muscles. (**C)** The psoas muscle can be stripped posteriorly-to-anteriorly by scraping out muscles attached to the anterior surface of the pelvis with a Penfield dissector. (**D)** Superior margin of the pelvic bone indicating the L6 vertebral body. (**E**) X-ray confirms the position of the pelvis relative to the L5/6 and L6/S1 intervertebral discs.

**Figure 2.**
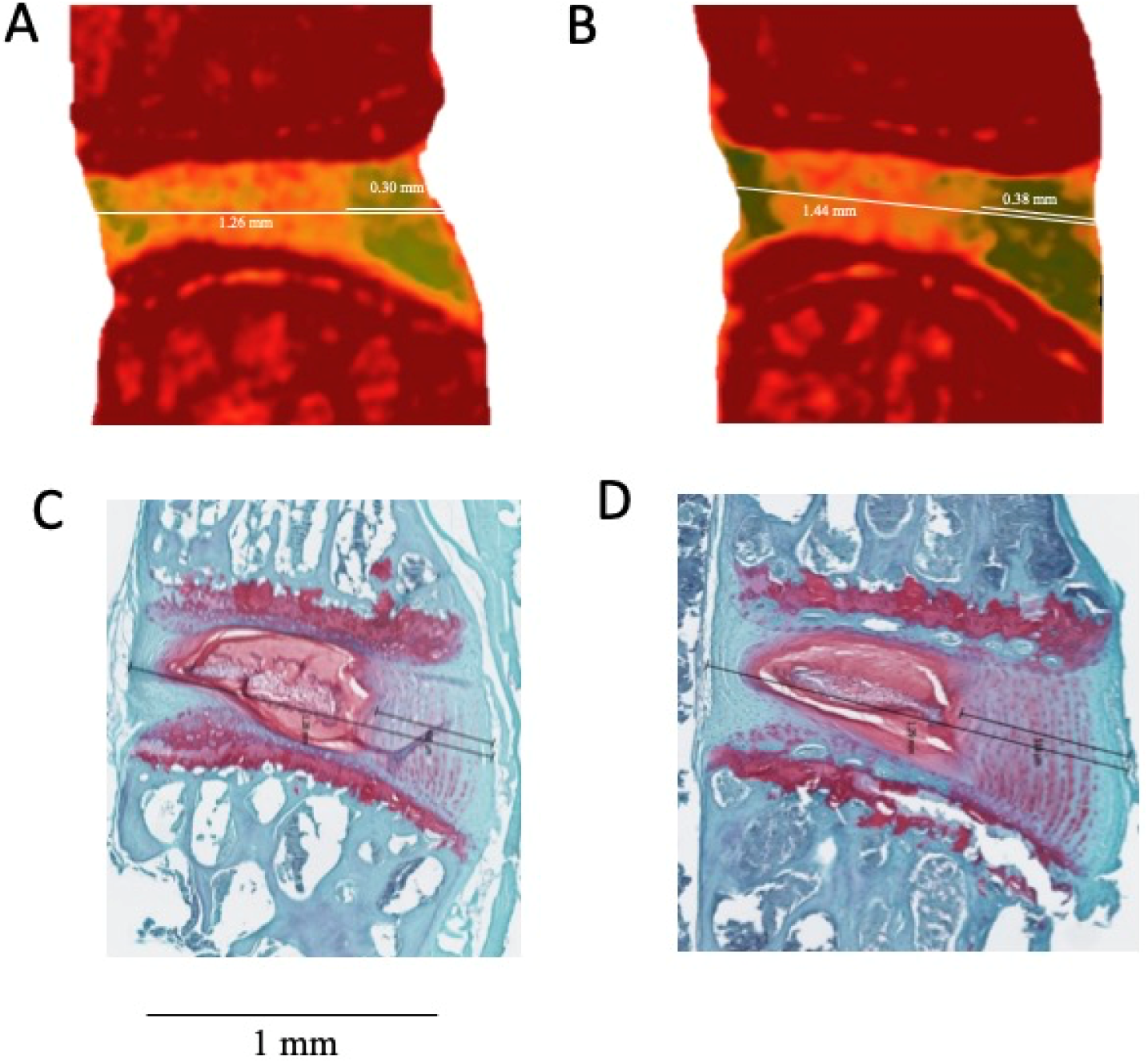
Measuring the thickness of the intervertebral disc and the annulus fibrosus **(A-B)** Contrast-enhanced microCT (CEμCT) and **(C-D)** histology were used to measure the IVD widths and anterior AF thicknesses. The CEμCT and histology measurements were highly correlated (r^2^ = 0.96, p < 0.001) and statistically indistinguishable (paired t-test, p = 0.37), confirming the fidelity of the non-destructive evaluation of CEμCT.

For the full-width injury, a 33G needle was inserted bilaterally through the IVD lateral axis of the IVD. In contrast to the partial-width injury, NP herniations were observed following the full-width injury.

### Contrast-enhanced microCT tomography (CEμCT)

Samples were incubated in a solution of 175 mg/mL solution diluted from a stock of OptiRay 350 (Guerbet, St. Louis) in PBS at 37°C. After 24 hrs of incubation, samples were scanned using a µCT40 (Scanco Medical, CH) at a 10-µm voxel size (45 kVp, 177 uA, high resolution, 300 ms integration).

CEµCT data was exported as a DICOM file for analysis in a custom MATLAB program. After an initial median filter (sigma = 0.8, support = 5), functional spine units were isolated from surrounding soft tissue not part of the IVD by drawing a contour around the outer edge every 5 transverse slices and morphing using linear interpolation. The IVD was manually segmented from the vertebral bodies with the same methodology as above. The remaining voxels were designated as the whole disc mask. The NP was thresholded from the AF followed by a morphological close and morphological open to fill interior holes and smooth the boundary. The volumes and average attenuations (intensity) were calculated from the mask of the NP and whole disc. The volume was determined from the total number of voxels contained within the mask and the attenuation was taken as the average 16-bit grayscale value of the voxels. Visualizations of the microCT were obtained using the image processing application OsiriX (Pixmeo, Geneva). AF thickness, partial-width injury depth and disc height index (DHI) were measured along the mid-sagittal plane. DHI was calculated as the ratio of the IVD height to width. IVD height was taken as the average at 5 equidistant points along the mid-sagittal plane. The ratio of NP intensity/disc intensity (NI/DI), defined as the average attenuation of voxels in the NP mask divided by the full disc mask, is an unbiased, fully three-dimensional measure that quantifies the relative size and hydration to inform the relative changes in degeneration ^18^. Samples where attenuation was saturated were not included in analysis.

### Histological preparation and evaluation

Following microCT, samples were fixed for 24 hours in 10% neutral buffered formalin followed by 3 days of decalcification in Immunocal (StatLab 1414-X). The samples were embedded in paraffin, sectioned at a thickness of 10 µm, and then stained with Safranin-O and Fast Green. Histological classification system recently developed by Melgoza et al. in 2021 was used to quantify the degeneration of the injured level (L6/S1) which allows the evaluation of the nucleus pulposus, annulus fibrosus, endplates and interface boundaries^19^. Morphology and NI/DI determined from CEµCT was used to further inform the level of degeneration of the injured level (L6/S1) compared to the internal control (L5/6).

### Measuring thickness of AF and the depth of partial-width injury

The CEμCT of the injured L6/S1 and histological analysis on uninjured L5/6 were used to measure the thickness of the AF. The CEμCT on the injured L6/S1 was used to measure the depth of injury as defined by the shortest perpendicular distance from the outer edge of annulus fibrosus to the visually observable outline of the injury site.

### Statistics

All statistics were examined for normality and nonparametric tests were used accordingly. A 2-way ANOVA was used to assess the effects of injury and postsurgical time on histological grade, attenuation and morphological parameters, with adjusted post hoc comparisons between groups (Tukey). Comparisons and effects were considered significant when the p-value is less than 0.05. All statistics were run using GraphPad Prism 9.3.1 (GraphPad, San Diego, California).

## Results

A total of 57 mice were subjected to the surgical procedure. The average surgery time was 15 min 38 sec ± 6 min 23 sec from incision to closure. There was no mortality or complications due to surgery. Caution was taken to prevent injury of the lumbosacral plexus located posteriorly during the blunt dissection of the psoas muscle from posterior to the anterior direction.

### Annulus Fibrosus (AF) Thickness

CEμCT and histology measurements of AF thickness and IVD width were highly consistent. The anteroposterior IVD width measured with CEμCT was 1.27 ± 0.13 mm (mean ± standard deviation), whereas that measured by histological analysis was 1.21 ± 0.11 mm. AF thickness measured with CEμCT was 0.38±0.05 mm and that measured with histological analysis was 0.43 ± 0.10 mm (Figure 3). The CEμCT measured values were statistically indistinguishable from (p = 0.37) and were highly correlated (r^2^ = 0.96) with the histologically measured values (Figure 2).

**Figure 3.**
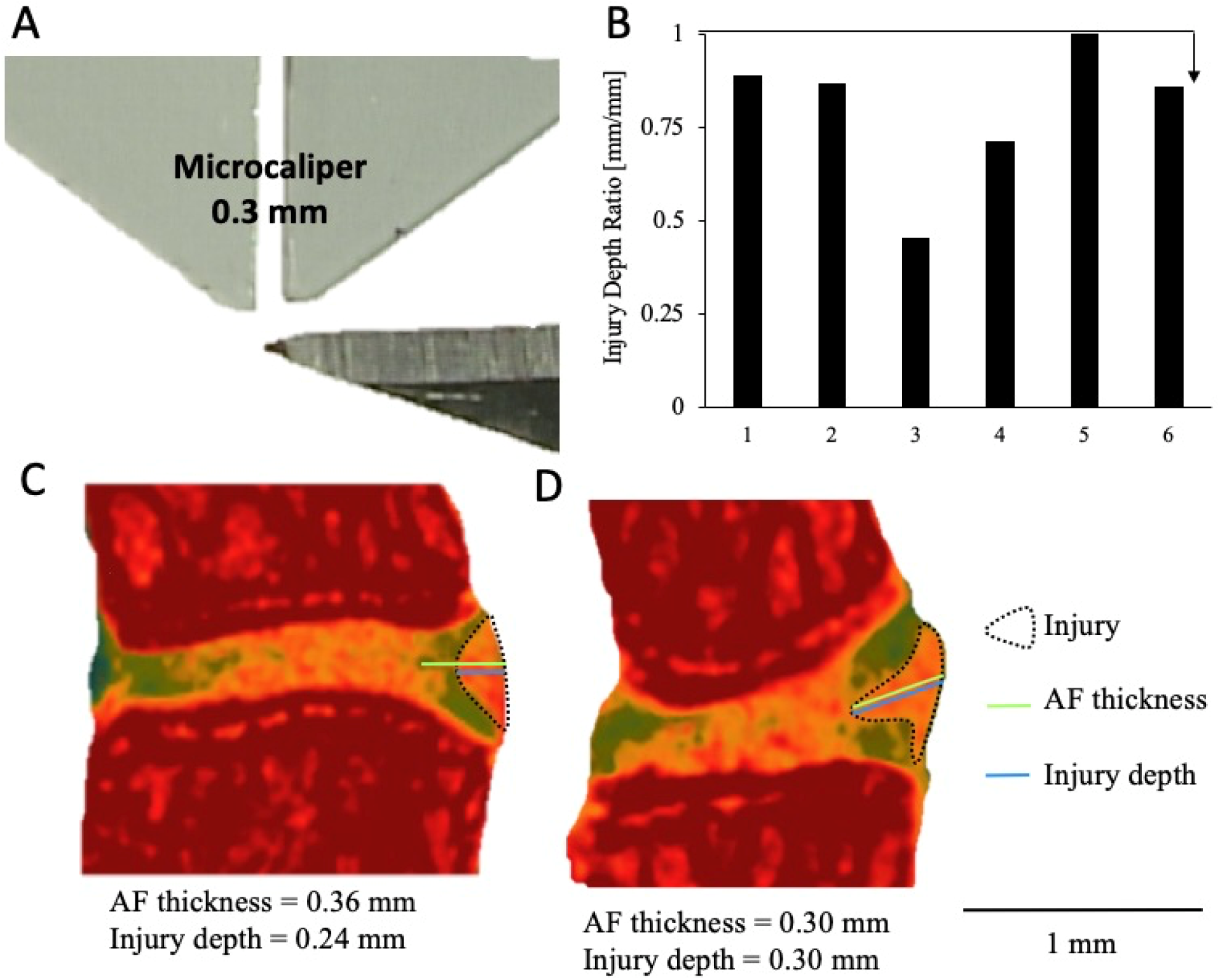
Depth of the partial-width injury measured by contrast-enhanced micro-computed tomography **(A)** The tip of the scalpel edge is marked at 0.3 mm to allow for visual confirmation during surgery that the partial-width injury is limited to the AF. **(B)** All of the injury depths were confirmed by CEµCT and histology to have depths that do not exceed of the AF in each injured IVD, confirmed by the ratio of injury depth to AF thickness which are less than 1 in all samples. **(C, D)** Representative CEµCT showing the range of depths achieved by this AF injury.

### Depth of Partial-width Injury

All injuries except one were isolated to the AF (Figure 3B). In sample 5, the NP may have been injured because CEµCT showed disruption of the AF/NP boundary (Figure 3C). The depth of injury measured with CEμCT was 0.29 ± 0.05 mm. The ratio of the injury depth to the AF thickness was 0.80 ± 0.19 (Figure 3D and 3E).

### Structural evaluation of the IVD

Neither the partial-width nor the full-width injuries resulted in changes in the Disc Height Index across the two-, four-, and eight-week time points (Figure 4D). Similarly, the size of the nucleus pulposus, characterized by the nucleus pulposus (NP) volume fraction, were not dramatically different between groups and time points (Figure 4E). The relative hydration of the NP, quantified by the ratio of NP attenuating intensity to that of the whole disc (NI/DI), revealed the full-width injury caused a dramatic decrease in the attenuation of the NP, indicating a loss of NP hydration, at the two- and four-week time points, with a trending change at eight-weeks.

**Figure 4.**
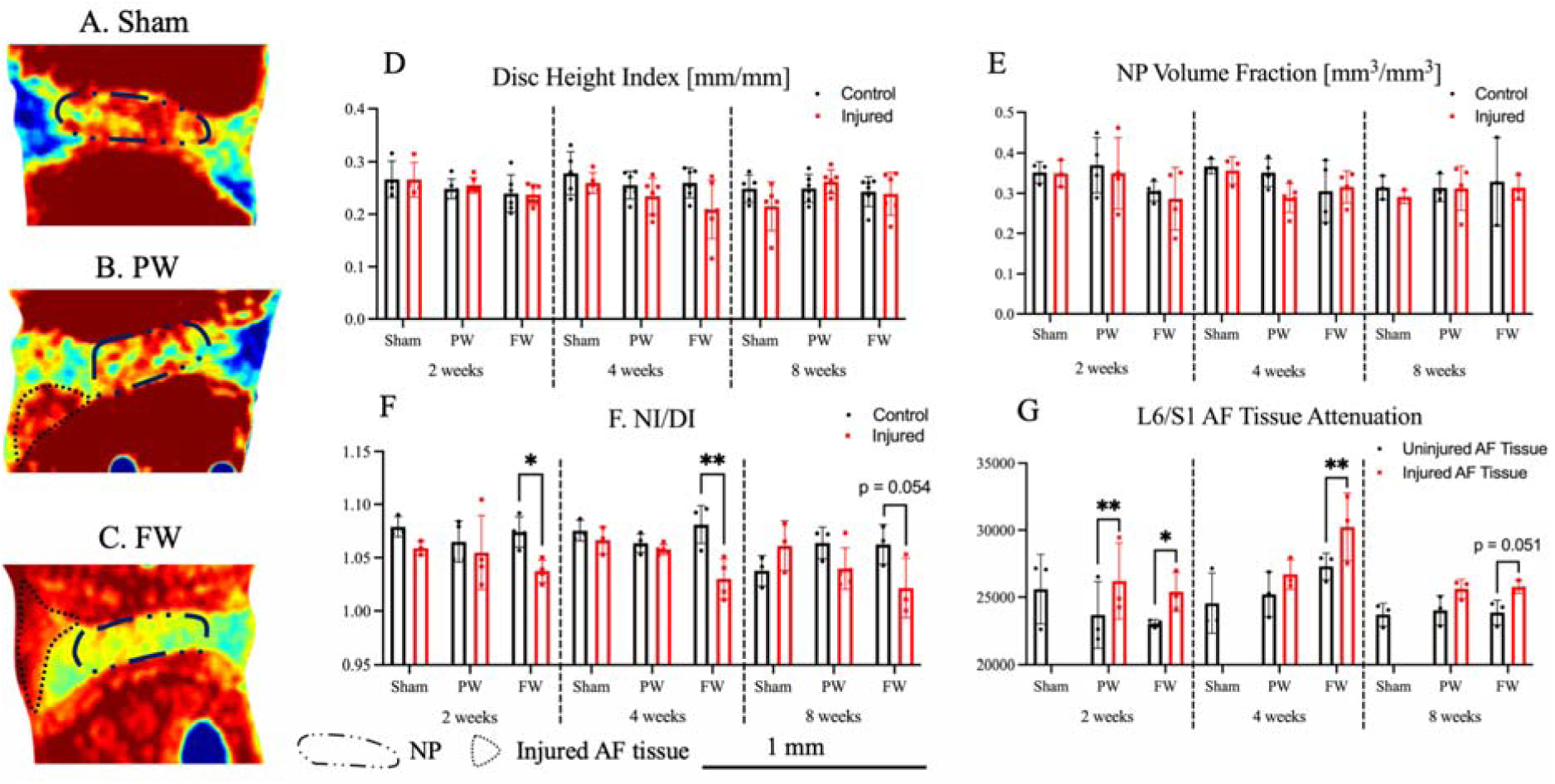
Full-width (FW) injury induces a sustained loss of NP hydration compared to both the partial-width (PW) injury and Sham groups. **(A-C)** Contrast-enhanced microCT of injured level (L6/S1) where injury site is visible in both FW and PW groups. **(D)** Mean attenuation of the AF in the injured and uninjured groups. Paired t-test indicates significant increase in attenuation at injury site compared to rest of uninjured AF (p < 10^−5^). **(E)** NI/DI of injured level (L6/S1) and uninjured control level (L5/6). While no changes in the whole IVD structure were detected between injured and uninjured levels with either FW or PW, the ratio of the nucleus pulposus intensity to the whole disc intensity (NI/DI – approximately 4% drop in intensity) revealed the loss hydration and proteoglycans in the nucleus pulposus of FW samples that is indicative of degeneration compared to the control IVDs.

The impact of the partial- and full-width injuries to the annulus fibrosus was observable at two weeks, The relative hydration of the annulus fibrosus (AF) increases after an injury, and this is evident in both the partial- and full- width groups. Two weeks after injury, both the partial- and full- injuries showed an increase attenuation of the AF, with the full-width injury group sustaining this increase (Figure 4G).

### Extent of IVD Degeneration

Quantitative histological analyses revealed a degenerative cascade to the IVD following full-width injury as early as two weeks post injury but not with the partial-width injury. The histological classification showed significant degeneration following FW injury at all timepoints (p < 0.05) in the NP, AF, interface boundaries, and total IVD score but not endplate score compared to Sham while no differences were detected between PW and Sham (Figure 5). No morphological differences between timepoints or injury groups were observed.

**Figure 5.**
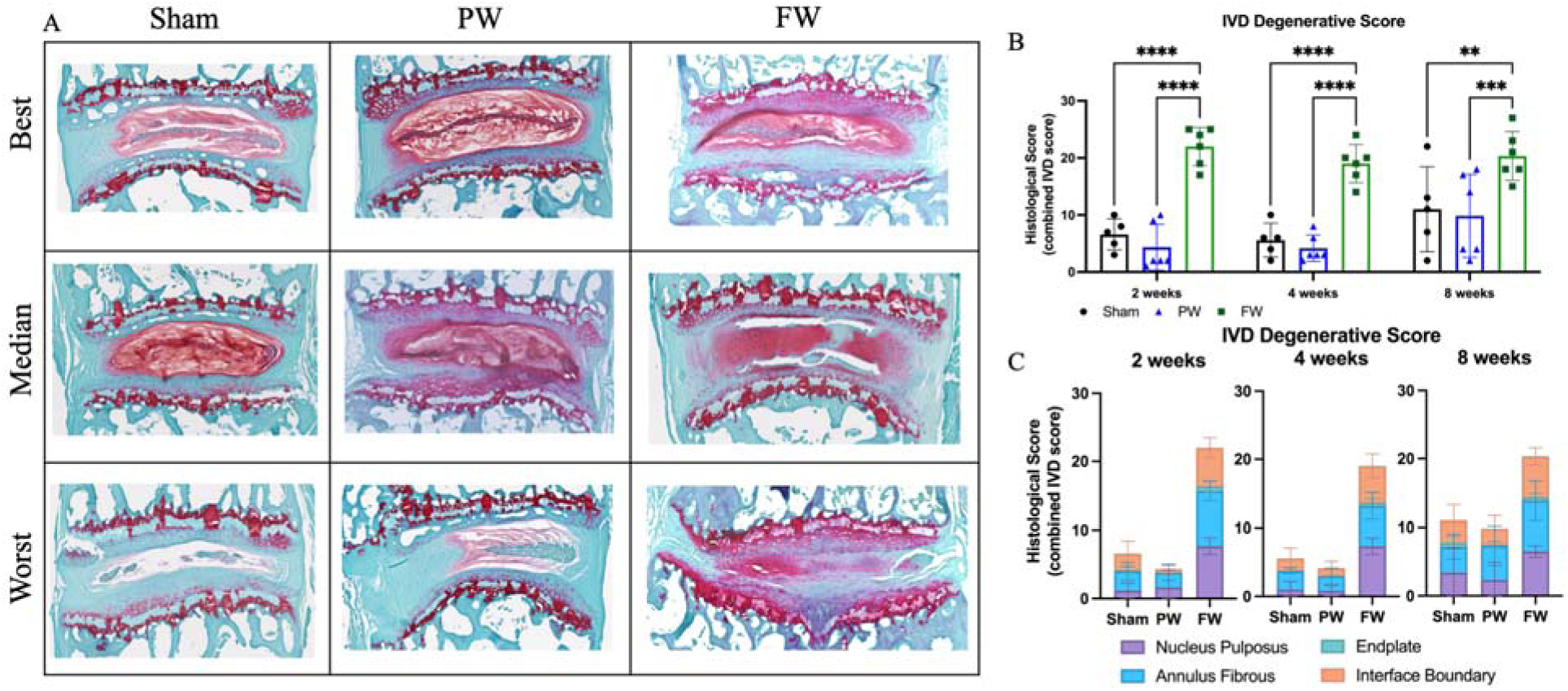
FW induces rapid and sustained IVD degeneration compared to both Sham and PW. **(A)** Safranin-O stained histological section of injured level (L6/S1) **(B)** Total IVD degenerative score of L6/S1. **(C)** Breakdown of NP, AF, endplate and interface boundary contributions to total IVD degenerative score. For both the PW and FW injuries, the site of injury were not observable on the histological sections after the 2-, 4- and 8-week post surgery. Significant degeneration was observed at all timepoints indicated by significant increases of degenerative scores in NP, AF, interface boundary and combined IVD degenerative grades in the FW group compared to Sham and PW (p < 0.05). While PW causes localized disruption to the AF, it does not appear to cause degeneration.

## Discussion

Animal models of IVD degeneration are imperative to elucidate the key molecular mechanisms of the pathophysiology. To date, rodents, rabbits, dogs, goats, sheep, and primates have been used as models for intervertebral disc degeneration^20^. One main advantage of mouse models is the availability of reagents and modifications that could be used concomitantly with surgically induced IVD degeneration. However, the small size of the mouse requires a high degree of surgical precision, particularly with access and exposure of the lumbar spine. While many studies utilize the mouse tail for degenerative models^12,21– 23^, the lumbar spine maintains the anatomical proximity to physiologically relevant structures such as the dorsal root ganglions.

We describe here a novel procedure for mouse lumbosacral IVD injury with visual guidance via microscopy and gross features. The retroperitoneal space of the spine can be located (Figure S1) with the left pelvic bone with gluteus muscles and proximal thigh as landmarks (Figure 1C and 1D). Blunt-dissection of the fat pad surrounding the thoracic cage anteriorly, the gluteus muscles superiorly, and the thigh muscles posteriorly (Figure 1B) exposes these two landmarks (Figure 1C). The lateral approach to the IVD allows for precise injury to the annulus fibrosus of the IVD (Figure 1). Exposure of the pelvic bone from the gluteus muscle and rotation of the pelvic bone posteriorly is necessary to achieve ample exposure of multiple IVDs in the surgical field. The psoas muscles can be easily scraped posterior-anteriorly with a Penfield dissector. CEµCT allowed for a 3D spatially unbiased quantification of morphology and composition^18^ that confirmed the localization of injury across the cross-section time points. CEµCT measurements of the IVD morphology were in excellent agreement with histological measures. NI/DI and histological analysis indicated degeneration of injured IVDs starting 2 weeks post-surgery.

The present study describes a surgical procedure to expose the lumbosacral spine of mice and to induce a localized experimental injury with AF-limited depth and compared the degenerative response with a more commonly used needle puncture injury. The technique described herein also focused on achieving a consistent depth and size of injury. The marked tip of the No. 11-scalpel blade tip was confirmed with CEμCT to provide AF-limited injury and resulted in no NP leakage under observation by microscopy. While most injury models cause damage to the nucleus pulposus, the cartilaginous endplates, and even the vertebral body^24–26^, the partial-width injury here results in an isolated injury to the outer annulus fibrosus. Surprisingly, we observed no degenerative changes in the endplate in the full-width injury, suggesting that it is possible to induce an IVD only injury using a carefully applied 33G needle. Following full-width injury, IVDs exhibited rapid degeneration as early as two weeks after injury, and this is sustained through the four- and eight-week time points. Despite maintaining disc height, the FW injury caused consistent degenerative changes in the nucleus pulposus, annulus fibrosus, and interface boundaries which included the AF-endplate and NP-AF boundaries. The PW injury did not produce statistically significant degenerative changes, but whether the innervation or vascularization changes at the outer annulus fibrosus remains to be investigated.

CEµCT allowed for nondestructive visualization of the injury site following both partial- and full-width injuries. The increased attenuation on CEµCT may indicate changes in composition and diffusion properties resulting from the disruption in the annulus fibrosus with both injuries. The injured annulus fibrosus appears to be highly hydrated, likely due to the increased interstitial fluids that localize around the injury site (Figures 3-4). Since IVD degeneration in humans often starts as a tear in the annulus fibrosus^17^, this model will enable mechanistic investigations of how injuries of the annulus fibrosus contribute to IVD degeneration and the subsequent development of low back pain. In contrast, the full-width injury to the nucleus pulposus induces a more rapid and severe course of degeneration that is more aligned with an acute trauma ^7,20,21^. Unlike the increased attenuation of the injured annulus fibrosus, the depressurized nucleus pulposus loses water and attenuates less than the healthy state (Figures 4). Consistent with unilateral injuries of the tail IVD, the resultant degenerative cascade was nuanced^11^, and required high resolutions modalities to detect measurable changes. Future studies will evaluate the partial-width injury as a slowly progressing IVD degeneration model following a clinically relevant injury.

## Supporting information

Video depicting surgery.

## Acknowledgements

This work is in part supported by NIH R01AR074441, R01AR077678, K01AR069116, P30 AR007992, and T32 DK108742. All authors contributed in the design, execution, and collection of data; as well as writing and critical review of the manuscript.

## Supplemental data

### Instrumentation

Instruments used in surgery included a size No. 11-scalpel blade (Henry Schein Medical), forceps (Fisher Scientific), dissecting scissors (VWR), Penfield dissector (VWR), q-tips (Dukal Corporation), 4-0 nylon sutures (Ethicon), suture grabbers (Fisher Scientific), and microscope (Zeiss).

### Methodology

*Retroperitoneal approach to the intervertebral disc procedure:* The left flank was shaved from the ventral to the dorsal midlines using veterinary trimmers. Mice were then transferred to the operating suite and positioned in the left lateral decubitus position. The skin was prepared for aseptic surgery via washing/rinsing with povidone iodine scrub and alcohol rinse. The animals were draped to isolate the prepped area. Isoflurane by inhalation was used to maintain surgical depth anesthesia throughout the procedure.

There were two surface landmarks utilized for this approach: 1) the anterior margin of the thigh (vertical) and bony prominence of the pelvis (horizontal) and 2) the fat pad reflected underneath the skin. The fat pad located between the anterior margin of the thigh and abdominal wall was utilized as an access point to the retroperitoneal space. A 1–1.5-cm horizontal incision was made along the bony prominence of the pelvic bone line covering the entry and the anterior 1/3 of the thigh (Figure S1A). The vertical caudal margin of the access point was retracted and resected by blunt dissection with Mosquito forceps (Figure S1B and S1C). The gluteus muscles and left pelvic bone (a) were utilized as a grip to hold and move the spine axis with forceps. The opening to the retroperitoneal space (c) between (a) and oblique abdominal muscles (b) was carefully blunt-dissected until the psoas muscle was exposed (Figure S1D).

**Supplemental Figure 1.**
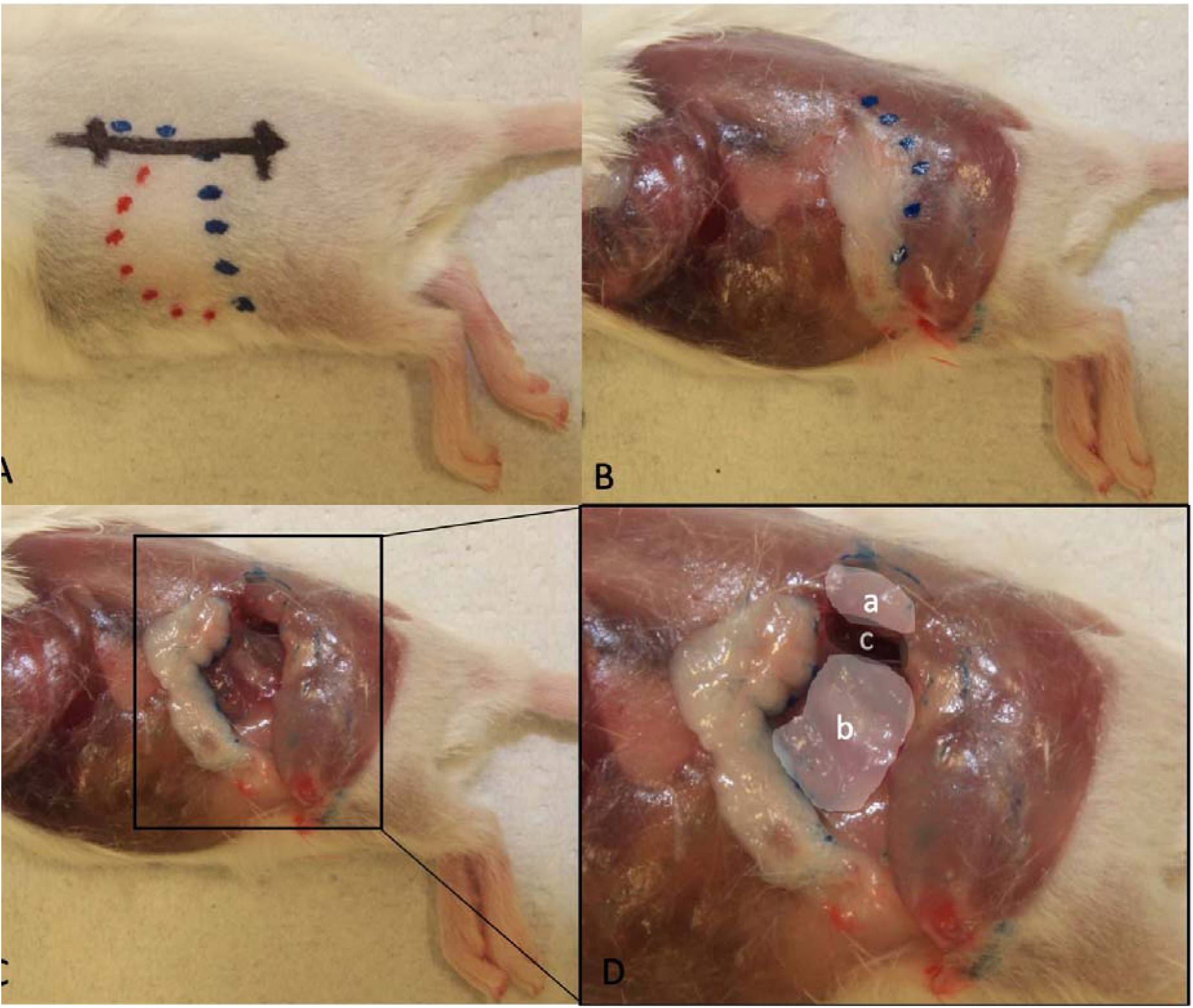
Lateral retroperitoneal approach to the mouse lumbosacral spine **(A):** The superficial anatomical landmarks are denoted by 1) the blue dotted line which outlines anterior margin of the thigh and bony prominence of the pelvis, and 2) the red dotted line which outlines the fat pad located between the thigh and abdominal wall. The black solid line is an approximately 1 cm incision along the pelvis line that reveals the surgical access point adjacent to the anterior thigh. (B and C): The access point to the retroperitoneal space lies between the blue and red dotted lines. The underlying fat pad and anterior margin of the thigh can be easily separated and retracted to expose the retroperitoneal space by blunt dissection. (**D):** The pelvis is rotated posteriorly to expose a broad working space by gripping (a) with forceps. The area (c) between (a) and (b) is carefully separated by blunt dissection to expose the psoas muscle: (a) gluteus muscles and left pelvic bone; (b) oblique abdominal muscles; (c) opening to the retroperitoneal space.

